# Infrared fiber optic spectroscopy detects bovine articular cartilage degeneration

**DOI:** 10.1101/2020.05.18.101600

**Authors:** Vesa Virtanen, Ervin Nippolainen, Rubina Shaikh, Isaac Afara, Juha Töyräs, Johanne Solheim, Valeria Tafintseva, Boris Zimmermann, Achim Kohler, Simo Saarakkala, Lassi Rieppo

**Affiliations:** Research Unit of Medical Imaging, Physics and Technology, Faculty of Medicine, University of Oulu, Oulu, Finland; Department of Applied Physics, University of Eastern Finland, Kuopio, Finland; Diagnostic Imaging Center, Kuopio University Hospital, Kuopio, Finland; School of Information Technology and Electrical Engineering, The University of Queensland, Brisbane, Australia; Biospectroscopy and Data Modeling Group, Faculty of Science and Technology, Norwegian University of Life Sciences, Ås, Norway

**Keywords:** infrared spectroscopy, attenuated total reflectance, articular cartilage, osteoarthritis

## Abstract

Articular cartilage (AC) is a soft connective tissue that covers the ends of articulating bones. Joint injuries may lead to degeneration of cartilage tissue and initiate development of post-traumatic osteoarthritis (OA). Arthroscopic surgeries can be used to treat joint injuries, but arthroscopic evaluation of cartilage quality is subjective. Therefore, new methods are needed for objective assessment of cartilage degeneration. Fourier transform infrared (FTIR) spectroscopy can be used to assess tissue composition based on the fundamental molecular vibrations. When combined with fiber optics and attenuated total reflectance (ATR) crystal, the measurements can be done flexibly without any sample processing. We hypothesize that Fourier transform infrared attenuated total reflection (FTIR-ATR) spectroscopy can detect enzymatically and mechanically induced changes similar to changes occurring during progression of OA. Fresh bovine patellar cartilage plugs (*n* = 60) were extracted and degraded enzymatically and mechanically. Adjacent untreated control samples (*n* = 60) were utilized as controls. Enzymatic degradation was implemented by 90-min and 24-hour collagenase as well as 30-min trypsin treatments. Mechanical damage was induced by: 1) dropping a weight impactor on the cartilage plugs, and 2) abrading the cartilage surface with a rotating sandpaper. Fiber optic FTIR-ATR spectroscopic measurements were conducted for control and degraded samples, and spectral changes were assessed with random forest (RF), partial least squares discriminant analysis (PLS-DA), and support vector machine (SVM) classifiers. RF (accuracy 93.1 % to 79.2 %), PLS-DA (accuracy 95.8% to 81.9%), and SVM (accuracy 91.7% to 80.6%) all had excellent classification performance for detecting the different enzymatic and mechanical damage on cartilage matrix. The results suggest that fiber optic FTIR-ATR spectroscopy is a viable way to detect minor degeneration of AC.

## 1 Introduction

Articular cartilage (AC) is a soft connective tissue that covers the ends of articulating bones. AC consists of extracellular matrix (ECM), chondrocytes and interstitial fluid. Main molecular components of ECM are collagens and proteoglycans along with lower amounts of glycoproteins and non-collagenous proteins (Sophia Fox *et al* 2009). The function of AC is to provide a low friction surface for smooth articulation of joints and redistribute the loads applied to the ends of bones. Osteoarthritis (OA), the most common musculoskeletal disease in the world, degenerates articular cartilage within synovial joints. OA is most common in knee, hip and hand joints. Since OA is the leading cause of disability among elderly people, affecting 250 million people worldwide, its impact on healthcare costs is significant (Hunter & Bierma-Zeinstra 2019).

Cartilage degeneration and changes in chondrocyte function are the most evident signs of progression of the disease (Pritzker *et al* 2006). In the early stages of the disease, the loss of proteoglycans, tissue fibrillation and degradation are seen at the superficial layer of AC. Excessive proliferation or hypertrophy of chondrocytes and cell death also occur due to OA. Furthermore, as OA progresses, chondron organization starts to disintegrate, and formation of vertical fissures is observed, making the AC surface discontinuous. In the latter stages of OA, the cartilage matrix erodes, and the bone is exposed. Thereafter, bone deformation and remodeling start to occur (Pritzker *et al* 2006). The etiology of OA is not completely known, and often OA is idiopathic, *i.e.*, it is not possible to identify a specific cause for the disease. One exception is post-traumatic OA, which is initiated by a trauma to the joint, *e.g.*, a sports injury or an accident. Early stage diagnostics in these cases is important, since successful intervention in the early stages of the disease can significantly postpone the progression of the disease (Lespasio *et al* 2017).

Arthroscopic surgeries are used to treat various knee OA symptoms, but the benefits of current methods are shown to be limited (Brignardello-Petersen *et al* 2017). The current arthroscopic classification of cartilage damage, which is based on manual mechanical probing of cartilage combined with visual inspection, is subjective. Experts have stated a need for improvement in differentiation of tissue damage (Spahn *et al* 2009). This subjective assessment may limit the outcomes of the arthroscopic operations due to misclassification of tissue health, and thus, there is a need for objective tools to assess tissue quality in order to guide surgical repair of joint injuries and reveal incipient post-traumatic OA (Favero *et al* 2015, Mandl 2019).

Several methods are currently under investigation for providing objective measures for surgeons. Intra-articular ultrasound (Virén *et al* 2009) and optical coherence tomography (OCT) (Li *et al* 2005) may be used to visualize the tissue structure to reveal *e.g.* the lesion depth. In addition to structural information, near-infrared (NIR) spectroscopy (Sarin *et al* 2018) can be used to assess the molecular composition of AC by studying the overtone frequencies of molecular vibrations. NIR light can penetrate through the whole cartilage tissue (Padalkar & Pleshko 2015), but the overtone vibrations are seen as broad features in NIR spectra, and the molecular sensitivity of NIR spectroscopy is limited (Bunaciu *et al* 2015).

Mid-infrared (MIR) spectroscopy provides better molecular selectivity than NIR spectroscopy as it excites the fundamental molecular vibrations. MIR region is routinely used in Fourier transform infrared (FTIR) microscopy to measure ECM molecule distributions in histological cartilage sections (Camacho *et al* 2001, Rieppo *et al* 2017). Furthermore, an FTIR spectrometer, combined with fiber optics and an attenuated total reflection (ATR) crystal, allows also flexible measurements of cartilage without any sample processing (West *et al* 2004, Hanifi *et al* 2013). In principle, these measurements could be done *in vivo* to provide objective information about the tissue properties, *e.g.*, during arthroscopic surgery. MIR frequencies can only penetrate a few micrometers through the sample surface (Raichlin & Katzir 2008), but the increased molecular selectivity of the MIR range may be advantageous especially when evaluating early OA changes.

We hypothesize that fiber optic FTIR-ATR spectroscopy can detect degenerative changes in AC associated with early OA. To test the hypothesis, we used fiber optic FTIR-ATR spectroscopy to differentiate mechanically and enzymatically damaged bovine cartilage from intact bovine cartilage. In enzymatic damage models, the collagenase treatment modeled superficial collagenase network damage, while trypsin treatment modeled proteoglycan depletion seen in the early stages of OA. Additionally, a longer collagenase treatment was used to induce major collagen network degradation occurring in the later stages of OA. In mechanical damage models, impact damage modeled tissue damage associated with high impact injuries, and surface abrasion simulated fibrillation seen on the surface of AC. FTIR-ATR spectra between intact and damaged tissue were compared to reveal the spectroscopic changes related to tissue damage. Furthermore, binary random forest (RF), partial least squares discriminant analysis (PLS-DA), and support vector machine (SVM) classifiers were trained to separate degenerated and untreated tissues based on their FTIR-ATR spectra.

## 2 Materials and Methods

### 2.1 Sample preparation protocol

Bovine (age 14 – 22 months) patellae (*n* = 12), obtained from a local slaughterhouse (Atria Oyj, Finland) were used in this study. Cylindrical (*d* = 7 mm) osteochondral samples were prepared from six different anatomical locations of each patella. The samples were divided into enzymatically degraded groups (*n* = 36), and mechanically degraded groups (*n* = 24). Control samples (*n* = 60) were prepared from locations adjacent to damage group samples as shown in Figure 1.

**Figure 1.**
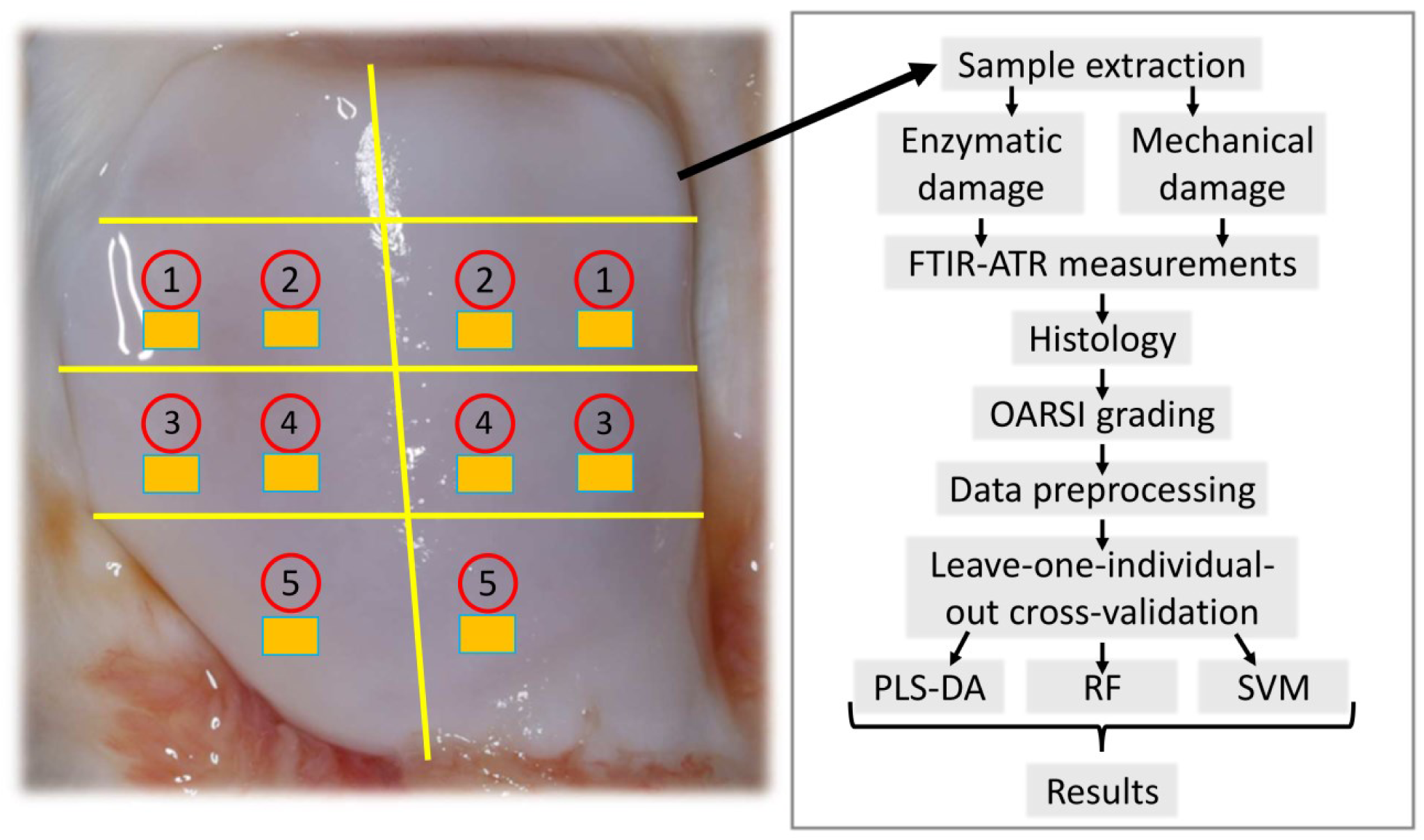
Sample extraction locations and complete flow diagram of the measurement protocol. Sample preparation locations of fresh bovine osteochondral plugs (*d* = 7 mm) are indicated in numbered red circles and adjacent control samples for each group are indicated with yellow rectangles. Numbers indicate following groups: 1) Collagenase 24-h, 2) Collagenase 30-min, 3) Trypsin 30-min, 4) Surface abrasion, 5) Impact loading.

Enzymatic degradation was targeted to the two main components of cartilage ECM: collagen network with collagenase D (0.29 U/mg, Sigma Aldrich) and proteoglycans with trypsin (T 4174, Sigma Aldrich). The samples were divided into 24-h collagenase (*n* = 12), 90-min collagenase (*n* = 12), and 30-min trypsin (*n* = 12) treated groups. Each sample was incubated at 37°C and 5% CO_2_ in phosphate buffered saline solution (PBS) containing the specified enzymes (0.1 mg/ml for collagenase D, and 0.5 mg/ml for trypsin) with added Penicillin-Streptomycin-Amphotericin B (100 units/ml Penicillin, 100 µg/ml Streptomycin and 0.25 µg/ml Amphotericin B, stabilized, Sigma-Aldrich). To prevent lateral penetration of the enzymes, enzymatic treatments were performed on larger rectangular pieces of tissue (10 - 15 mm per side), and afterwards cylindrical plugs (*d* = 7mm) were extracted from the selected locations.

Two types of mechanically induced damage were applied in this study: impact loading (*n* = 12) and abrasion (*n* = 12). Impact loading was conducted using a stainless-steel impactor (*m* = 200 g) with a polished steel ball (*d* = 1 cm) acting as the impact surface. The impactor was dropped from a height of 7.5 cm. This height was chosen based on preliminary testing to create cracks on the surface of AC. Surface abrasion was induced using a custom-made tool enabling 180° rotation of a metal plate with P80 sandpaper (particle size 200 µm) attached to the surface of the plate. A constant stress of 4 kPa was applied to the cartilage surface during two 180° rotations.

All samples were measured within 10 hours of extraction from the knee joint. In between treatments and measurements, the samples were stored in PBS solution with protease inhibitors to minimize the effects of natural degradation of fresh tissue. After the mechanical and enzymatic treatments, all samples were allowed to recover in PBS for a duration of 60 minutes to facilitate the diffusion of damaged tissue components. All samples were kept in 4°C to reduce natural degradation of the tissue until measurements for the specific sample plug occurred.

### 2.2 Sample measurement protocols

FTIR spectroscopic measurements were conducted utilizing a custom ATR probe (Art Photonics GmbH, Germany) that was connected to a Thermo Nicolet iS50 FTIR spectrometer (Thermo Nicolet Corporation, Madison, WI, USA), equipped with a Globar mid-infrared source and a liquid nitrogen-cooled mercury cadmium telluride (MCT) detector. The samples were measured 3 times, with a new probe contact established for each measurement, resulting in 3 technical replicates for each sample. Measurements were done with 2 cm^-1^ spectral resolution, digital spacing of 0.2411 cm^-1^, and averaging 64 scans over the range from 4000 cm^-1^ to 400 cm^-1^. The background spectrum was measured for each sample separately. Measurements were controlled by OMNIC software (Thermo Nicolet Corporation, Madison, WI, USA).

### 2.3 Histology

After the experiments, the sample plugs were fixed in formalin, decalcified in ethylenediaminetetraacetic acid (EDTA), and embedded in paraffin. Subsequently, 3-µm-thick histological sections were cut and stained using Safranin-O for qualitative evaluation of the cartilage damage.

### 2.4 Data analysis

Preprocessing by multiplicative signal correction (MSC) (Martens & Stark, 1991) was applied, and spectral range was truncated to 1800-900 cm-1. To compare the spectra of treated and control groups, mean spectra of each damage and control group were calculated. Difference spectra were calculated by subtracting the mean damage group spectrum from the mean control spectrum of each group (Figure 3a). Furthermore, second derivative of difference spectra were calculated using Savitzky-Golay algorithm using a window size of 91 (corresponding to 22 wavenumbers) to emphasize subtle differences (Figure 3b).

**Figure 2.**
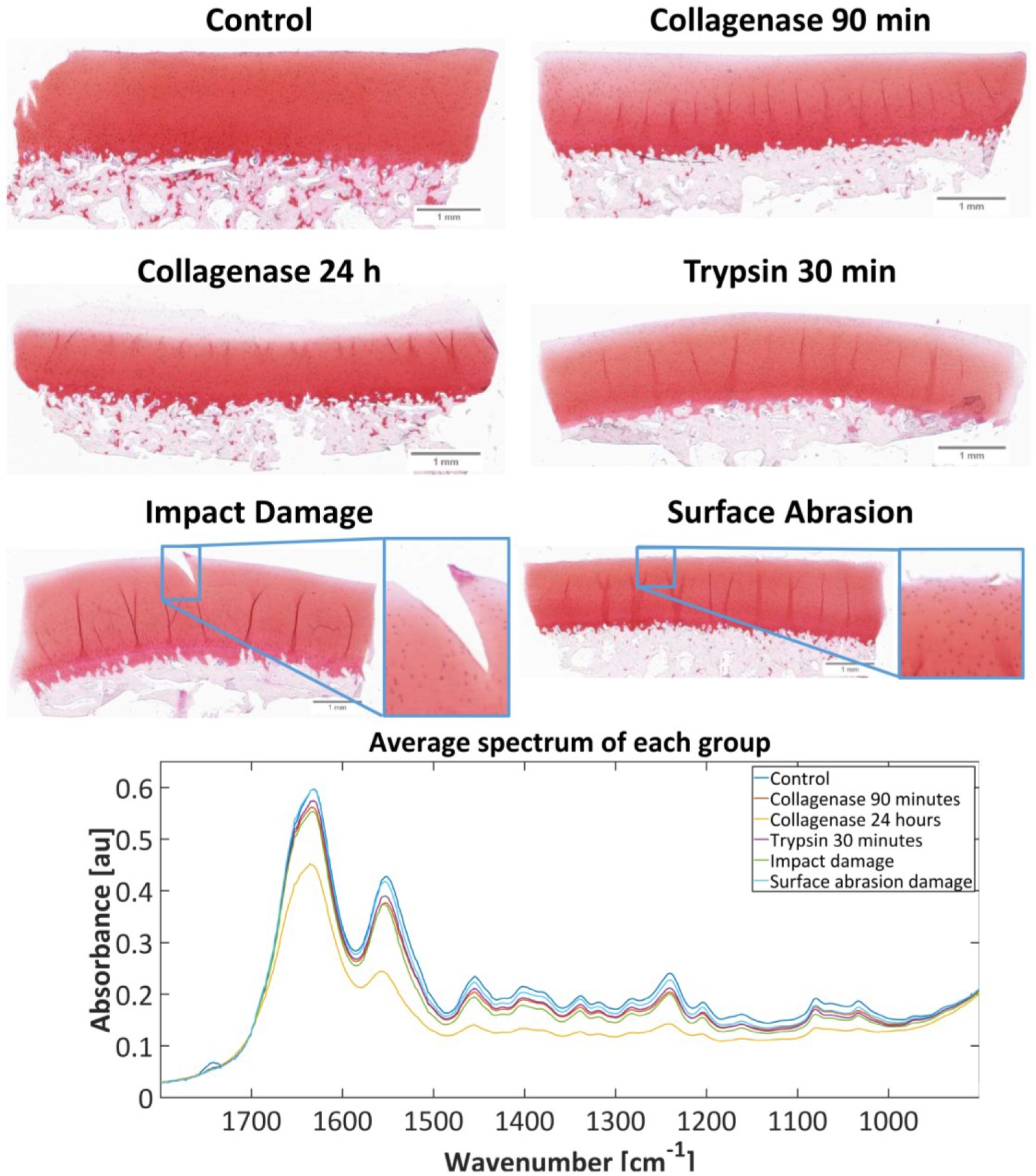
Safranin-O stained histological sections visualizing the damage induced by different treatments (top); mean spectra of the damage groups and the control group (bottom).

**Figure 3.**
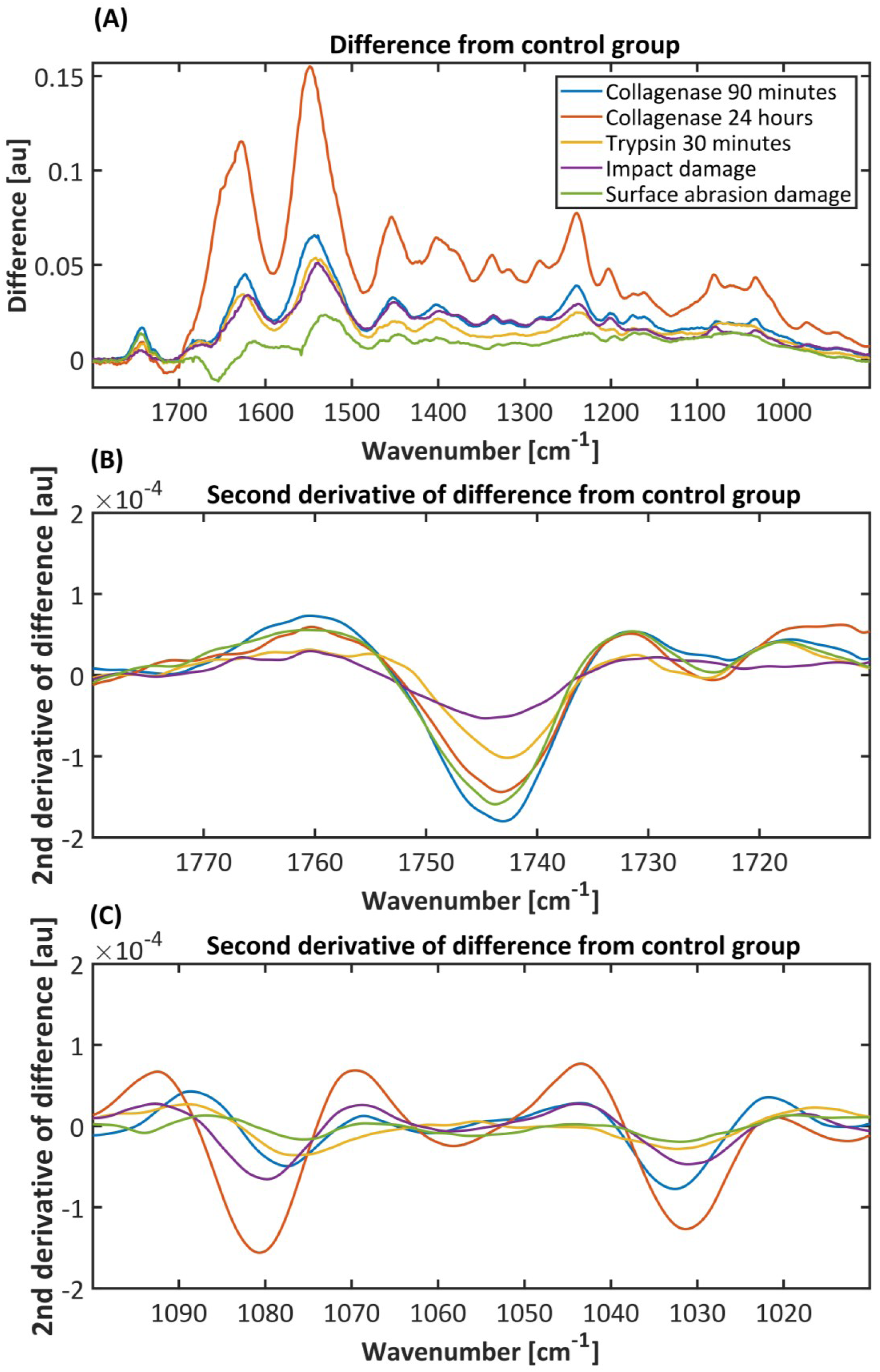
a) Difference spectrum of each group (mean of control group spectra subtracted from mean of damaged sample spectra for each group); b) Second derivative spectrum of each group’s difference spectrum from the region 1780-1710 cm^-1^. c) Second derivative spectrum of each group’s difference spectrum from the region 1100-1010 cm^-1^.

RF, PLS-DA, SVM classifiers were used for classifying the spectra into treated and untreated groups. Binary models were built by utilizing the three technical replicates for each of the damage group samples and the adjacent control samples (sample locations are illustrated in Figure 1).

RFs utilize multiple individual decision trees and combine the outputs into a final prediction using majority vote (Ho, 1995). The main advantage of RF classifier is that it is robust to noise and overfitting, and it is well suited for data with large number of features, such as spectroscopic data. The number of RF trees was set to 100. In PLS-DA, independent (*i.e.* spectra) and dependent (*i.e.* sample groups) variables are used to calculate latent variables so that the covariance between the two variable sets is maximized. The dimensionality of the data is being reduced thus reducing the multi-collinearity problem. To avoid overfitting of the models, the number of PLS components was selected by checking the minimum classification error of models built using 1-7 PLS components and selecting the smallest number of components where the error is within 5% of the true minimum. The number of PLS components chosen for each model is shown in the Table 2. SVMs utilize mapping of training data into a new hyperspace with the help of a kernel function, after which it constructs an optimal hyperplane that fits the data best (Brereton & Lloyd, 2010). SVMs perform well in fitting non-linear data. For SVM classifiers, Bayesian optimization was used to find the optimal gamma and C values for the radial basis function kernel.

**Table 1.** Assignments of notable molecular vibrations observed in difference spectra between damage and control groups. The features are seen in difference spectra, second derivative spectra, as well as PLS-DA regression coefficients shown in Figures 3-5. (Fung *et al* 1998, Holman, *et al* 2003, Belbachir *et al* 2009, Zelig *et al* 2011, Vidal & Mello 2016, Rieppo *et al* 2017, Talari *et al* 2017)

| Wavenumber (cm<sup>-1</sup>) | Assignment |
| --- | --- |
| 1743 | Ester carbonyl C=O stretching (phospholipids) |
| 1720-1580 | Amide I band; C=O & C-N stretching |
| 1580-1490 | Amide II band; N-H bending, C-N & C-C stretching |
| 1458 | CH<sub>3</sub> asymmetric bending (Collagen) |
| 1403 | CH<sub>3</sub> symmetric bending (Collagen) |
| 1338 | CH<sub>2</sub> side-chain vibrations (Collagen type II integrity) |
| 1317 | CH<sub>2</sub> stretching (Collagen) |
| 1300-1200 | Amide III band |
| 1283 | Collagen |
| 1238 | Collagen |
| 1203 | Collagen type I |
| 1180-1000 | Carbohydrate region (C-O, C-O-H, C-O-C) |
| 1165 | Collagen type I |
| 1080 | C-O stretching of carbohydrate residues (Collagen) |
| 1034 | C-O stretching of carbohydrate residues (Collagen) |

**Table 2.** Performance of RF, PLS-DA, and SVM binary classifier models. Accuracy (ACC), sensitivity (TPR), and specificity (TNR) are shown for all methods. Best (or shared best) performance of classifiers indicated in bold. Number of PLS-DA components utilized for each PLS model shown in column “PLS comps”.

|  | *Random Forest* |  |  | *PLS-DA* |  |  |  | *SVM* |  |  |
| --- | --- | --- | --- | --- | --- | --- | --- | --- | --- | --- |
|  | ACC | TPR | TNR | ACC | TPR | TNR | PLS comps | ACC | TPR | TNR |
| *Enzymatic damage* |  |  |  |  |  |  |  |  |  |  |
| Collagenase 24 h | **91.7 %** | 91.7 % | **91.7 %** | 90.3 % | **100 %** | 80.5 % | 1 | **91.7 %** | 91.7 % | **91.7 %** |
| Collagenase 90 min | 90.3 % | **91.7 %** | 88.9 % | **95.8 %** | **91.7 %** | **100 %** | 3 | 80.6 % | 75.0 % | 86.1 % |
| Trypsin 30 min | 84.7 % | 77.8 % | **91.7 %** | 81.9 % | 72.2 % | **91.7 %** | 2 | **86.1 %** | **86.1 %** | 86.1 % |
| *Mechanical damage* |  |  |  |  |  |  |  |  |  |  |
| Impact damage | 79.2 % | 75.0 % | 83.3 % | **95.8 %** | **91.7 %** | **100 %** | 2 | 91.7 % | **91.7 %** | 91.7 % |
| Surface abrasion | **93.1 %** | **94.4 %** | **93.1 %** | 90.3 % | 88.9 % | 91.7 % | 3 | 84.7 % | 77.8 % | 91.6 % |

Leave-one-individual-out cross-validation was used to estimate the performance of the established models. Cross-validation error is usually an overoptimistic performance estimation of a model. Calculating the error of prediction by taking out all the spectra of one individual (with all replicates) at each step of the cross-validation ensures that no samples of that individual were used in establishing the model. This type of cross-validation is the closest to real external validation error and should be used when no external test set validation is possible due to limited sample size.

Spectral data processing and data analyses were performed by algorithms developed in house in the environment of MATLAB, R2018a (The Mathworks Inc., Natick, MA, United States).

## 3 Results

Representative Safranin-O stained sections along with the mean spectrum of damage and control samples for each group are shown in Figure 2. As expected, the largest absolute change occurred in collagenase 24-h group, where most of the surface layer is missing completely due to major collagen network disintegration. The collagen network disruption in both collagenase groups led also to significant proteoglycan depletion at the superficial layer of AC. In contrast, trypsin induced a significant proteoglycan loss while keeping the collagen network visibly intact. In mechanically induced damage groups, surface abrasion led to fibrillation of cartilage surface along with minor proteoglycan depletion, whereas impact damage resulted in deeper fissures along with proteoglycan loss.

The most notable differences between the damage and control groups in the FTIR-ATR spectral data were observed in Amide I, II and III regions (Table 1). Spectra of damaged cartilage showed decreased absorbance compared to controls in all cases. The greatest difference between the control and damage groups was seen in the 24-h collagenase group. In contrast, the surface abrasion group was observed to be the least different from the control. The differences between the damage and control groups can be seen from the difference spectra (Figure 3a) as well as from the second derivative of difference spectra (Figures 3b & 3c).

Multiple collagen peaks, *e.g.* at 1458 cm^-1^, 1403 cm^-1^, 1338 cm^-1^, 1317 cm^-1^, 1238 cm^-1^, 1203 cm^-1^, 1165 cm^-1^ and 1080 cm^-1^ (Table 1), can be identified from the difference spectra (Figure 3a). The peak at 1338 cm^-1^ corresponds to CH_2_ side-chain vibration of collagen and it is more specifically related to the integrity of type II collagen. The peak at 1317 cm^-1^, originating from CH_2_ stretching, has also been shown to be associated with collagen (Fung *et al* 1998). These peaks were clearly seen in the difference spectra of both collagenase groups and impact damage group (Figure 3a), while trypsin and surface abrasion damage groups did not show such changes. Similarly, the collagen C-O stretching peaks at 1080 cm^-1^ and 1034 cm^-1^ were only seen in the difference spectra of collagenase and impact damage groups.

RF models had accuracies between 93.1 % and 79.2 %, while PLS-DA accuracies ranged from 95.8% to 81.9%, and SVM accuracies from 91.7% to 80.6%. Sensitivity (TPR) for RF classifiers ranged from 94.4 % to 77.8%, PLS-DA classifiers from 100 % to 72.2 %, and SVM classifiers from 91.7 % to 75 %. Specificity (TNR) for RF classifiers ranged from 93.1 % to 88.9 %, PLS-DA from 100 % to 80.5 % and SVM from 86.1 % to 91.7 % (Table 2).

Regression coefficients of five different PLS-DA models are shown together in Figure 4. Substantial contribution of Amide I and Amide II regions to the classification between damaged and control samples was seen in all treatment groups. PLS-DA regression coefficients for all groups show contribution from the lipid peak at 1743cm^-1^, which is especially visible in the surface abrasion group, while showing only minor contribution to the model in 24-h collagenase group. Furthermore, carbohydrate region shows complex contribution for classification in all groups. For collagenase treatments and impact damage groups, the most important additional feature was Amide III band.

**Figure 4.**
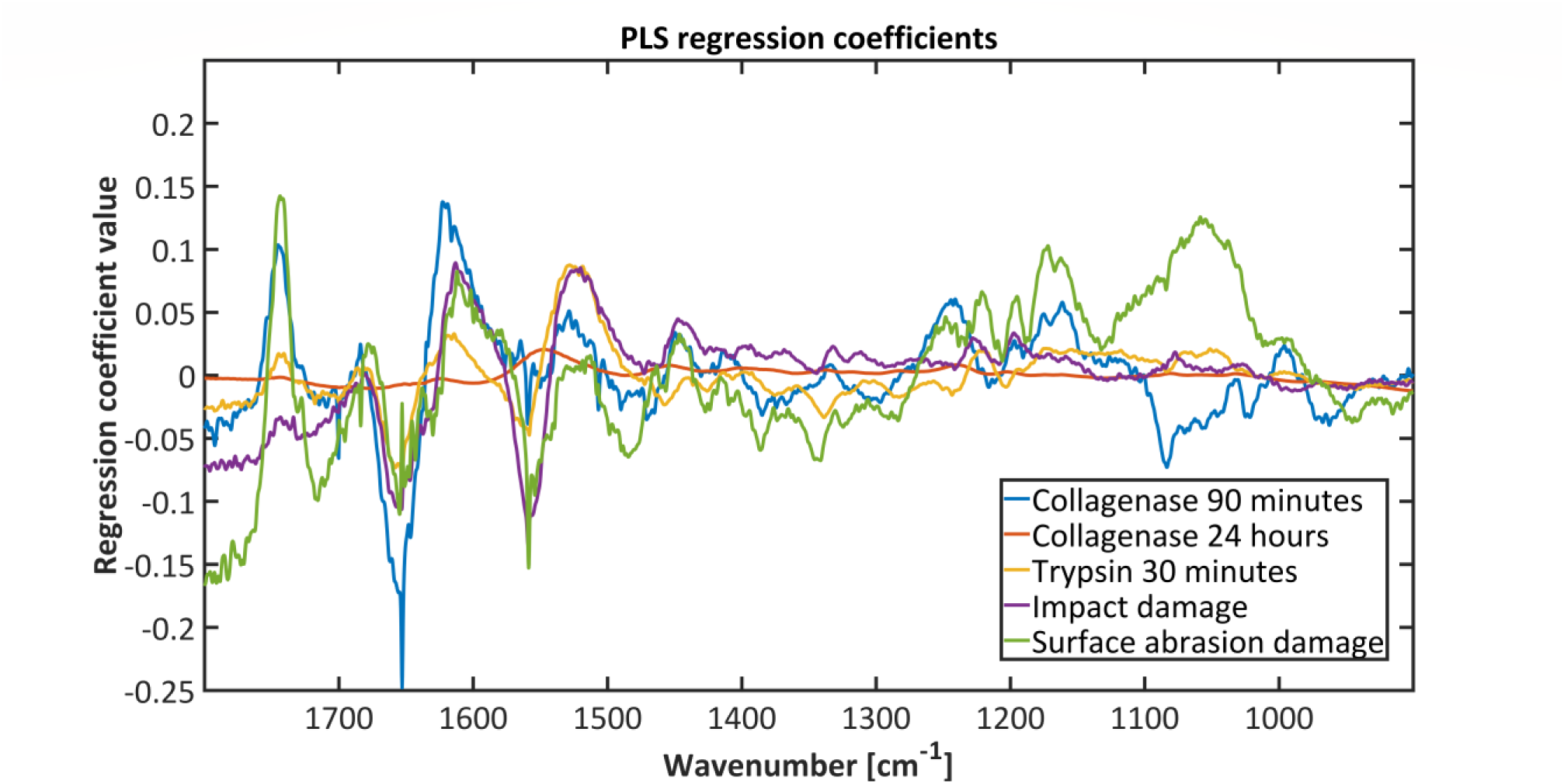
Regression coefficients for five PLS-DA models established separately for each damage group.

## 4 Discussion

In this study we hypothesized that fiber optic FTIR-ATR spectroscopy can detect degenerative changes in AC associated with early OA. Bovine AC specimen were damaged either mechanically or enzymatically to simulate early stage (collagenase 90-min & trypsin 30-min), post-traumatic (impact damage & surface abrasion), and late stage (collagenase 24-h) OA. The results of this study show that FTIR-ATR spectroscopy combined with multivariate classification methods, such as RF, PLS-DA, and SVM, can distinguish degenerated AC from intact normal tissue with good classification accuracy, sensitivity, and specificity. The enzymatic and mechanical damage models were observed to create degenerative changes similar to those occurring in OA. The findings encourage further investigation of more complex OA models accounting for other tissue level changes occurring during the development and progression of OA.

FTIR spectroscopic imaging has been previously reported to be well suited for cartilage tissue measurements (Camacho *et al* 2001), and it has also been used to measure enzymatically degraded cartilage (Potter *et al* 2001, West *et al* 2005, Rieppo *et al* 2012). However, to our knowledge, this is the first time when enzymatically and mechanically damaged fresh bovine cartilage has been studied with FTIR fiber optic probe. Bovine patellar cartilage has been shown to be suitable to be used in models of early human OA since it closely resembles human cartilage in thickness, morphometry and zonality (Hargrave-Thomas *et al* 2013). The use of fresh cartilage allowed a controlled spectroscopic comparison of intact and degenerated tissues, and the results suggest that fiber optic FTIR-ATR spectroscopy could also be suitable for *in vivo* cartilage measurements. In earlier ex vivo studies, fiber optic FTIR-ATR spectra have been shown to correlate to histopathological OA grade in human and rabbit cartilage (Hanifi *et al* 2012, O’Brien *et al* 2015). The results of this study show that FTIR-ATR spectroscopy can also detect small degenerative changes, suggesting it may be useful for evaluating the presence of incipient post-traumatic OA e.g. after joint injuries.

Safranin-O staining is a sensitive histological method for analyzing the extent of proteoglycan loss in AC (Schmitz *et al* 2010). Based on Safranin-O staining, the trypsin treatment showed degradation of proteoglycans in superficial layer. Even though trypsin is often used to deplete cartilage proteoglycans, it has been shown that it is not completely specific to proteoglycans, since it also has a minor degradative effect on already cleaved collagen (Harris *et al* 1972). 90-min collagenase treatment showed depletion of proteoglycans as a result of collagen network disruption, while the surface layer structure was fully disintegrated in the 24-h collagenase treatment group. Impact damage group samples had large cracks extending to mid or deep zone. The compression from the impact damage most likely caused depletion of proteoglycans due to collagen network disintegration. Slight proteoglycan loss can be seen on the histological Safranin-O stained sections close to the edges of the cracks. Surface abrasion group shows fibrillation of the surface layer, while having limited effect on Safranin-O staining intensity, indicating the damage to the tissue is not extending beyond the surface.

Spectral changes are in line with the results observed in histology. Collagenase enzyme group had reduced intensity in the collagen peaks. 90-min treatment showed moderate and 24-hour treatment showed major changes. Somewhat similar spectral changes were observed in trypsin treatment group compared to collagenase degradation groups. One distinct difference, however, is that the changes to collagen peaks at 1337, 1080 cm^-1^ and 1034 cm^-1^ (Vidal & Mello, 2016) are not as pronounced in the trypsin or surface abrasion groups when compared to other damage groups. Surface abrasion and trypsin groups mainly showed spectral changes attributed to the loss of proteoglycans. On the other hand, the impact damage group showed collagen network disorganization, causing spectral changes which were more similar to collagenase treatment groups.

One interesting spectral feature that could be seen in all pre-treatment groups was a lipid peak at 1743 cm^-1^. This peak is not visible or is minor in the treated samples. Normal cartilage tissue contains a small amount of lipids (Villalvilla *et al* 2013). A surface-active phospholipid bilayer has been shown to be present in superficial layer of cartilage in normal synovial joints (Schwarz & Hills, 1998). One possible explanation for the lipid peak weakening is that the damage treatments may have exposed the lipids embedded within the articular surface and allowed them to dissolve in the PBS solution. The lack of surface-active lipid bilayer is seen on damaged AC surface (Beldowski *et al* 2017, Jung *et al* 2017). Furthermore, the EDTA present within PBS solution during storage could have caused the lipids to dissolve. This kind of effect of EDTA on lipids has been shown on artificial lipid membranes (Prachayasittikul *et al* 2007). It might be possible that the damage treatments have accelerated the effect of EDTA lipid destabilization in comparison to undamaged samples. Ability to differentiate this change within the surface of AC with fiber optic spectral measurements offers possibilities for future studies of phospholipid interactions within synovial joints and opens up possible development of further diagnostic tools.

The classifiers selected for this study (RF, PLS-DA, and SVM) have been used in similar tasks before (Kohler *et al* 2010, Hanifi *et al* 2013, Lee *et al* 2018, Tafintseva *et al* 2018, Wang *et al* 2019). The models established using these classifiers provided excellent classification on our dataset. PLS-DA achieved best accuracy for collagenase 90-min and impact damage groups, while RF and SVM achieved a shared best accuracy for collagenase 24-h group. RF had best accuracy also for the surface abrasion group, while SVM was able to classify trypsin damage most accurately. Overall, all classifiers were able to differentiate the treated cartilage from untreated cartilage. PLS-DA showed slightly more consistent performance when compared to RF and SVM. Furthermore, the PLS-DA models were relatively simple as they had only 1-3 PLS components. Therefore, it could be argued that the linear PLS-DA should be preferred over the non-linear models. However, non-linear models established by RF and SVM, may prove to be efficient for more heterogenous data, *e.g.*, when osteoarthritic human cartilage is studied.

One limitation of this study is the small number of samples, which is due to the sample measurement protocol. It is known that freezing cartilage tissue may change its biochemical and biomechanical properties (Qu *et al* 2014), and, therefore, the use of fresh cartilage is important especially when evaluating the suitability of the

method for *in vivo* measurements. Not being able to freeze the samples limited the number of samples which could be prepared and measured in a reasonable time. For this reason, an external test set could not be used to estimate classification model performance. However, leave-one-individual-out cross-validation is a good estimate of model’s accuracy in such case. A larger dataset is needed to establish more reliable and robust models and validate the obtained results.

Methods that are able to assess degeneration of cartilage, especially during early stages of OA, are actively developed. The potential of fiber optic FTIR-ATR spectroscopy for assessing OA grade of human cartilage has been demonstrated earlier (Hanifi *et al* 2012). In this study, we demonstrated that fiber optic FTIR-ATR spectroscopy is also sensitive to minor cartilage damage, which could be useful, *e.g.*, in knee arthroscopy after an injury. Objective measures of cartilage quality provided by fiber optic FTIR-ATR spectroscopy could improve the arthroscopic evaluation of cartilage damage, and potentially also improve the outcome of repair surgeries by providing accurate objective information of the extent of damaged tissue.

## 5 Conclusion

Fiber optic FTIR-ATR spectroscopy offers a viable way to detect mechanical and enzymatic degeneration of fresh AC. These degenerative changes simulate the pathological changes occurring during the progression of OA. PLS-DA, RF and SVM classifiers combined with FTIR-ATR spectroscopy may offer additional tools for objective assessment of AC health during arthroscopy in the future.

## Acknowledgements

This work was supported by the Europe Union’s Horizon 2020 research and innovation programme (H2020-ICT-2016-2017) project MIRACLE (grant agreement number 780598), and the Academy of Finland (project number 310466).

